# The inducible *lac* operator-repressor system is functional in zebrafish cells

**DOI:** 10.1101/2020.02.10.939223

**Authors:** Sierra S. Nishizaki, Torrin L. McDonald, Gregory A. Farnum, Monica J. Holmes, Melissa L. Drexel, Jessica A. Switzenberg, Alan P. Boyle

## Abstract

**Background:** Zebrafish are a foundational model organism for studying the spatio-temporal activity of genes and their regulatory sequences. A variety of approaches are currently available for editing genes and modifying gene expression in zebrafish, including RNAi, Cre/lox, and CRISPR-Cas9. However, the *lac* operator-repressor system, a component of the *E. coli lac* operon which has been adapted for use in many other species and is a valuable, flexible tool for studying the inducible modulation of gene expression, has not previously been tested in zebrafish.

**Results:** Here we demonstrate that the *lac* operator-repressor system robustly decreases expression of firefly luciferase in cultured zebrafish fibroblast cells. Our work establishes the *lac* operator-repressor system as a promising tool for the manipulation of gene expression in whole zebrafish.

**Conclusions:** Our results lay the groundwork for the development of *lac-*based reporter assays in zebrafish, and adds to the tools available for investigating dynamic gene expression in embryogenesis. We believe that this work will catalyze the development of new reporter assay systems to investigate uncharacterized regulatory elements and their cell-type specific activities.

## Background

Experimental approaches for the study of transcriptional regulation by cis-regulatory elements *in vivo* require methods for both genetically modifying cells or organisms, and for measuring expression levels of specific genes. Zebrafish (*Danio rerio*) is an ideal model organism for investigating the spatio-temporal-specific regulation of gene expression throughout the developing embryo as it satisfies the requirements for ease of genetic manipulation and expression readout. Microinjection of DNA into fertilized embryos allows for simple and effective delivery of genome-modification tools, such as *Tol2* transposons, that mediate genomic integration of constructed expression cassettes. Additionally, the transparency of zebrafish embryos facilitates the observation of fluorescent signals from reporter genes within live cells and tissue. Due to its benefits as a model organism, many technologies for studying gene function have been developed in zebrafish, including Cre/lox [1], tamoxifen-inducible Cre [2], the Tet-On system [3], RNAi [4, 5], and more recently, CRISPR based-methods [6]. However, the use of the *lac* operator-repressor system, a tool which functions transiently in a native context with minimal disruption of local regulation compared to many of the aforementioned methods, has yet to be demonstrated in zebrafish.

The *lac* operator-repressor system is an inducible repression system established from studies of the *lac* operon in *Escherichia coli* (*E. coli*) that regulates lactose transport and metabolism [7]. The Lac repressor (LacI) binds specifically to a *lac* operator sequence (*lacO*), inhibiting the *lac* promoter and *lac* operon expression through steric hindrance [8]. Addition of the allosteric inhibitor Isopropyl β-d-1-thiogalactopyranoside (IPTG) to cells frees the *lac* operon to express its associated gene by inhibiting the binding of LacI to *lacO* sequences. The use of IPTG with the *lac* operator-repressor allows for inducible reversal of transcriptional repression.

Since its discovery in prokaryotes, the *lac* operator-repressor system has been modified for use in eukaryotic organisms to study the regulation of gene transcription [8–11]. Experiments in mammalian cell lines from mouse, monkey, and human [8–10, 12], as well as in whole mouse [13], demonstrate the utility of the *lac* operator-repressor system. It has also successfully been applied in cell lines and whole organisms of the amphibian axolotl, suggesting that this system can be utilized in a wide range of organisms [14]. Modifications to the *lac* operator-repressor system has allowed for constitutive, ubiquitous expression [8, 15, 16], visually assessed output [10, 12], and the ability to study both gene repression and activation [17, 18], emphasizing its flexibility for studying gene expression dynamics. The ability of IPTG to relieve repression in the *lac* system makes it a more adaptable tool for studying the temporal dynamics of gene expression, compared to constitutively active or repressed reporter gene systems.

In this paper, we provide evidence that the *lac* operator-repressor system can function in the zebrafish fibroblast cell line PAC2, adding a versatile new tool for the study of zebrafish genetics and transcriptional regulation. The results in a zebrafish cell line support the potential functionality of the *lac* operator-repressor system to function in whole zebrafish. In addition, we demonstrate that the CMV-SV40 enhancer-promoter produces strong, widespread reporter gene expression in both a zebrafish cell line and embryos. This enhancer-promoter combination provides a flexible, non-tissue specific expression module for zebrafish to aid in reporter gene detection at a cellular level.

## Results

### The CMV-SV40 enhancer-promoter shows widespread expression in zebrafish

In order to promote the repression of a reporter gene in our assay, we sought to increase LacI expression in transfected cells by including a strong enhancer-promoter driving LacI. The CMV enhancer and promoter are frequently used in reporter vector construction across a wide range of studies due to their strong and constitutive promotion of gene expression. This includes zebrafish where the CMV enhancer-promoter has previously been shown to have potent tissue-specific expression [19]. However, recent studies have demonstrated that the tissue-specific promoter function of non-CMV promoters may be lost when paired with the CMV enhancer [20, 21]. With the goal of identifying enhancer-promoter pairs driving non-tissue specific gene expression in whole zebrafish, we examined CMV enhancer activity paired with a non-CMV promoter, the SV40 minimal promoter. The function of the CMV-SV40 enhancer-promoter was first validated in PAC2 cells by inserting them upstream of luciferase in a pGL3 plasmid. Relative luciferase output of the CMV-enhanced SV40 pGL3 plasmid was compared to a pGL3 plasmid containing only a minimal SV40 promoter. The CMV-SV40 enhancer-promoter was able to drive a 24-fold increase in luciferase expression compared to the promoter-only control, suggesting the CMV-SV40 enhancer-promoter is able to function as a strong enhancer-promoter combination in PAC2 cells (**Fig. 1A**).

**Figure 1.**
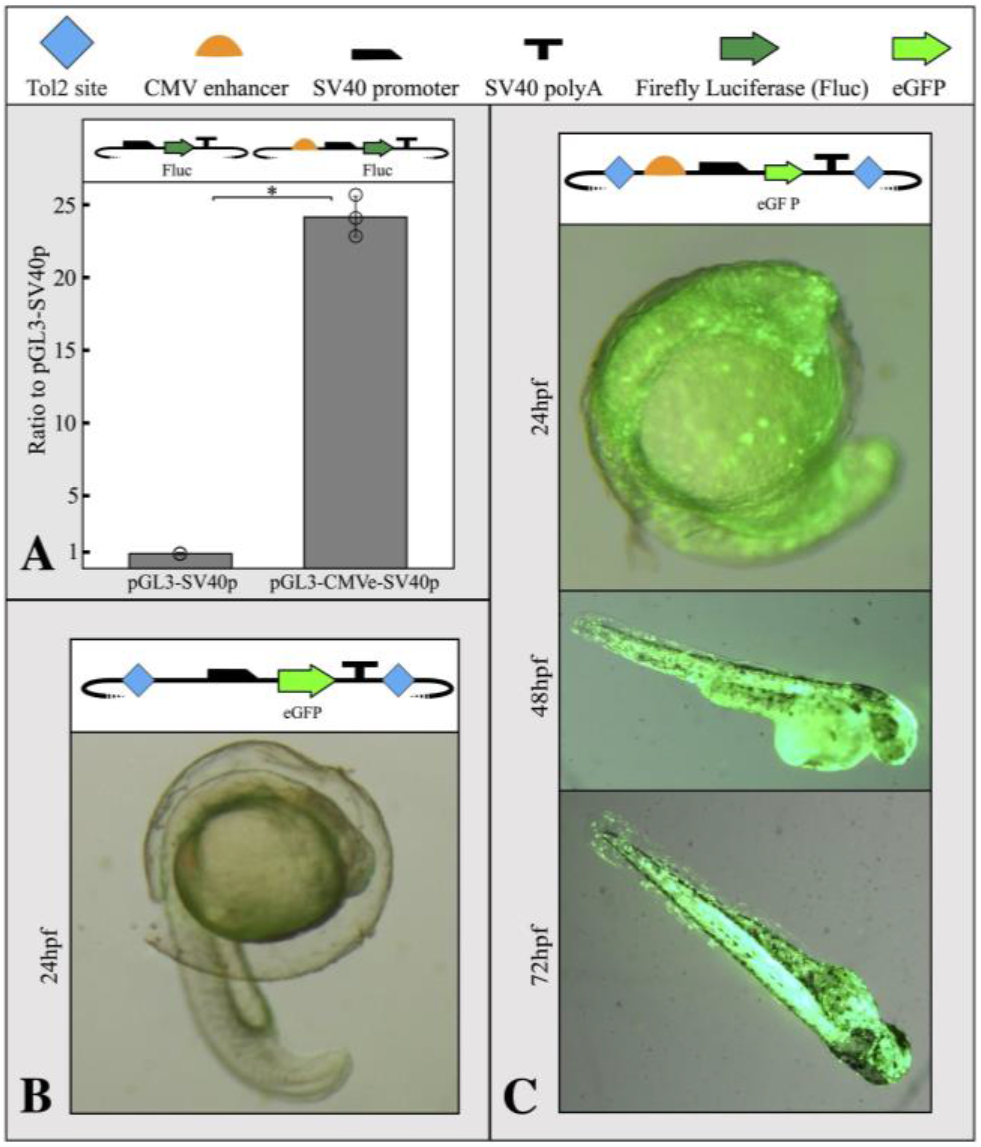
The CMV-SV40 enhancer-promoter drives widespread reporter gene expression in PAC2 cells and zebrafish embryos. A) CMV-SV40 enhancer-promoter shows a 24-fold increase in luciferase activity in PAC2 zebrafish fibroblast cells when transfected with a CMV-SV40 enhancer-promoter driving luciferase compared to a SV40-only plasmid. Error bars represent standard deviation of 3 biological replicates. Statistical significance determined using a stand 2-sided T-test * P-score < 0.01. Points represent values for all 3 replicates in each condition. B) SV40 promoter-only plasmids did not result in observable eGFP expression. C) CMV-SV40 enhancer-promoter driving eGFP shows non-tissue specific expression in zebrafish up to 72 hours post-fertilization. All plasmid components for each transfection design are detailed as symbols at the top of the figure. The TSS begins where the SV40 promoter begins to slope downward.

To determine if the CMV-SV40 enhancer-promoter was able to enhance reporter expression in whole zebrafish, the *Tol2* transposon system was utilized to integrate eGFP-expressing test plasmids into zebrafish embryos. As in PAC2 cells, two constructs were evaluated; one containing a CMV-SV40 enhancer-promoter upstream of an eGFP reporter gene, and one with only a minimal SV40 promoter. While no detectable level of eGFP activity in the promotor-only control was observed (**Fig. 1B**), composite brightfield and GFP images of 24, 48 and 72 hours post-fertilization embryos injected with the CMV-SV40 enhancer-promoter construct showed non-tissue specific eGFP expression (**Fig. 1C**).

### The *lac* operator-repressor system is functional in the PAC2 zebrafish cell line

To test the functionality of the *lac* operator-repressor system in zebrafish, a repressible reporter plasmid containing 6 *lac* operators in the 5’UTR of the firefly luciferase gene and a LacI-expressing plasmid were co-transfected into PAC2 cells. When a plasmid expressing a non-functional LacI (NFLacI) gene was co-transfected, no repression was observed (**Fig. 2**), whereas a plasmid expressing CMV enhancer-driven levels of LacI resulted in about 65% repression. An intermediate level of repression (∼40%) was observed when LacI was expressed from a plasmid containing only a SV40 minimal promoter, indicating that the extent of repression correlates with LacI levels in the cell. Addition of IPTG to the cells resulted in full relief of repression in all cases. This indicates that LacI is responsible for repression of luciferase expression in these cells.

**Figure 2.**
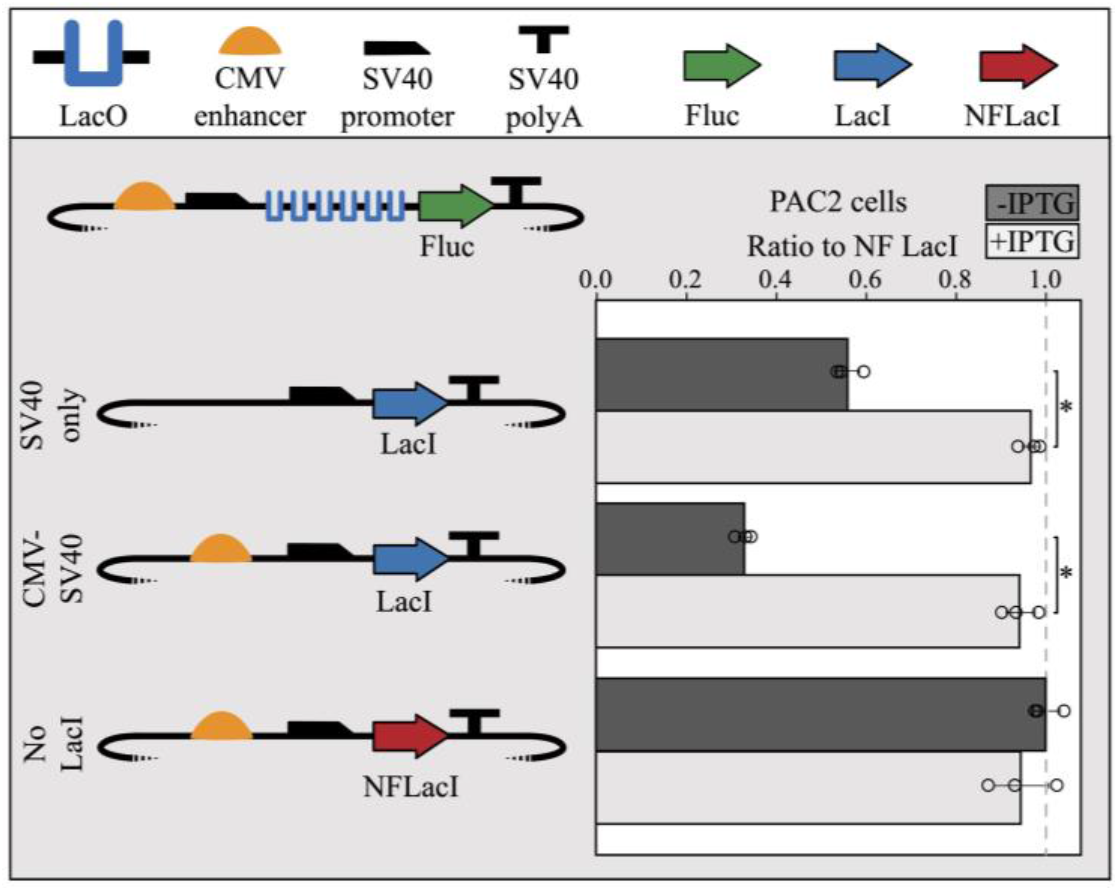
Co-transfection of LacI-expressing modules with repressible reporter modules result in LacI-mediated repression in PAC2 cells. SV40 promoter-only driven expression of LacI shows moderate repression (40%) and CMV-SV40 enhancer-promoter driven expression of LacI shows high repression (70%) of a repressible module containing 6x LacO sites. Expression of non-functional LacI (NFLacI, frameshift mutant) shows no repression and all modules showed maximal reporter expression in the presence of 1mM IPTG. Error bars represent standard deviation of replicates (n=3). Points represent values for all 3 replicates in each condition. The dashed line shows the NF LacI IPTG-negative control. The TSS begins where the SV40 promoter begins to slope downward. Statistical significance determined using a stand 2-sided T-test * P-score < 0.001.

To compare PAC2 repression levels to previously published data [22] and test broad functionality within different cell lines, we replicated the LacI experiment in the K562 human cell line. When both cell types were co-transfected with the same plasmid mixture, the performance of the *lac* operator-repressor system was nearly identical in PAC2 and K562 cells (**Sup. Fig. 2**). Both PAC2 and K562 cells showed around 60-65% repression when co-transfected with a molar equivalent of CMV enhancer-driven LacI containing plasmid (∼400ng), and roughly 10-20% repression when co-transfected with a similar molar equivalent of SV40 promoter-only driven LacI-expressing plasmid. These results demonstrate that the *lac* operator-repressor system functions in PAC2 cells at a comparable level to human K562 cells.

## Discussion

Zebrafish are a commonly used model organism for studying the spatio-temporal dynamics of *cis-*regulatory element activity and gene function. However, the flexible and widely used *lac* operator-repressor system has previously been untested in zebrafish. Here we demonstrate that the *lac* operator-repressor system functions in zebrafish cells, consistent with observed activity in other model eukaryotic systems.

For the development of a reporter system in whole organisms like zebrafish, it is critical to demonstrate non-tissue specific activity of an enhancer to provide robust output for single-cell reporter signal detection. The CMV enhancer is routinely used in reporter assays to drive strong and constitutive gene expression, and in a more widespread nature when paired with a non-CMV promoter [20, 21]. We provide quantitative evidence that the CMV-SV40 enhancer-promoter robustly increases luciferase gene expression over an SV40-only control plasmid in zebrafish fibroblasts. Furthermore, strong, non-tissue specific eGFP signal was observed in fertilized zebrafish eggs that persisted 72 hours post-fertilization after integration of a CMV enhancer controlled SV40-eGFP reporter construct. These robust levels of expression are critical in whole-organism studies where only a small number of cells may be expressing a reporter gene, and a high level of expression from a non-tissue specific enhancer-promoter, such as CMV-SV40, may facilitate their detection.

Changes in expression of the repressor protein LacI are inversely related to changes in reporter expression. This response appears to provide a level of repression directly related to the LacI level, rather than functioning as an on/off switch. This will allow for a more nuanced measure of *lac* regulatory control. Upon the addition of IPTG, luciferase signal was recovered to the level of a non-functional LacI control, indicating that robust repression is completely reversible at low IPTG concentrations. The pronounced response to IPTG treatment, as well as minimal toxicity in a zebrafish cell line, suggest the *lac* operator-repressor system is a viable tool for use in whole zebrafish.

*Lac* operator-repressor systems can be used to control endogenous gene expression without interrupting native regulatory processes, as *lacO* sites can be inserted in benign regions, such as introns and UTRs. Transcriptional inhibition of RNA polymerase by steric hindrance can achieve repression without introducing artificial modifications to the locus and causing prolonged alterations in regulatory behavior. This is in contrast to other systems that achieve transcriptional control by tethering a protein domain with activating or silencing effects through chromatin modifying or other endogenous mechanisms. Specificity of repression is also less of a concern compared to novel CRISPRi methods known for off-target effects [23]. As demonstrated by the REMOTE-control system, the lac system can also be used in conjunction with Tet-related systems to drive both activation and repression of a single loci, bringing additional flexibility to zebrafish studies [17]. This system allows for time-controlled experiments, where a reporter gene is repressed only for a limited time window, making it a crucial tool for replicating the restriction of gene expression during development.

## Methods

### Plasmid design

CMV-SV40 enhancer-promoter luciferase plasmids were generated by restriction digestion to insert a CMV enhancer and a minimal SV40 promoter, or only a minimal SV40 promoter, upstream of a luciferase reporter molecule in the context of a pGL3 plasmid (Promega, E1751). Plasmids designed for whole zebrafish injection were generated by replacing the firefly luciferase reporter with an enhanced green fluorescent protein (eGFP) reporter, and adding flanking minimal *Tol2* 200 base pair 5’ sequence and 150 base pair 3’ sequence for integration into the genome [24]. The sequence of the CMV-SV40 promoter is as follows: GGCATTGATTATTGACTAGTTATTAATAGTAATCAATTACGGGGTCATTAGTTCATAGCCCATATATGGAGTTCCGCGTTACATAACTTACGGTAAATGGCCCGCCTGGCTGACCGCCCAACGACCCCCGCCCATTGACGTCAATAATGACGTATGTTCCCATAGTAACGCCAATAGGGACTTTCCATTGACGTCAATGGGTGGAGTATTTACGGTAAACTGCCCACTTGGCAGTACATCAAGTGTATCATATGCCAAGTCCGCCCCCTATTGACGTCAATGACGGTAAATGGCCCGCCTGGCATTATGCCCAGTACATGACCTTACGGGACTTTCCTACTTGGCAGTACATCTACGTATTAGTCATCGCTATTACCATGGACTTGCATCTCAATTAGTCAGCAACCATAGTCCCGCCCCTAACTCCGCCCATCCCGCCCCTAACTCCGCCCAGTTCCGCCCATTCTCCGCCCCATGGCTGACTAATTTTTTTTATTTATGCAGAGGCCGAGGCCGCCTCTGCCTCTGAGCTATTCCAGAAGTAGTGAGGAGGCTTTTTTGGAGGCCTAGGCTTTTGCAAAAAGCTC.

*Lac* operator-repressor system plasmids were created using the EMMA golden gate assembly method [25]. All plasmids assembled using EMMA have a backbone consisting of an ampicillin resistance gene and a high-copy-number ColE1/pMB1/pBR322/pUC origin of replication. Backbone elements are denoted by terminating dotted lines in all plasmid schematics (**Fig. 1, Fig. 2, Sup. Fig. 1, Sup. Fig. 2**). The EMMA toolkit was a gift from Yizhi Cai (Addgene kit # 1000000119) [25]. The *lacI* CDS and C-terminal NLS were cloned from the Addgene plasmid pKG215 and inserted into an EMMA entry vector to create an EMMA part. pKG215 was a gift from Iain Cheeseman (Addgene plasmid # 45110) [26]. A frameshift mutation was introduced by inserting an adenosine in the fourth codon of *lacI* to create a non-functional LacI (NFLacI) for use in control experiments. The LacI-expressing module contains a minimal SV40 promoter, the *lacI* gene, and a SV40 polyA tail, with or without the addition of an upstream CMV enhancer (**Fig. 2**). The repressible reporter plasmid includes a CMV enhancer and a minimal SV40 promoter upstream of a firefly luciferase gene with symmetric *lac* operators inserted in its 5’UTR, terminated by a SV40 polyA tail. In order to maximize repression activity, six copies of the *lac* operators containing the sequence AATTGTGAGCGCTCACAATT were utilized in this study. This sequence is the “symmetric” *lac* operator that possesses tighter binding with LacI than the canonical *lac* operator sequences [17].

### Cell culture

The zebrafish fibroblast cell line PAC2 was maintained as previously reported [27]. Cells were grown at 28 degrees Celsius in Leibovitz’s L-15 +glutamine Medium (Invitrogen, 21083027) containing 15% heat inactivated fetal bovine serum (FBS; Sigma-Aldrich, F4135-500ML) and 1% antibiotic-antimycotic (Corning, MT30004CI) until confluent. Confluent cells were washed with 1x phosphate buffered saline (PBS; Invitrogen, 10010023) and detached from the plate with 0.05% Trypsin-EDTA for 5 minutes (Invitrogen, 25300054). Trypsin was quenched with FBS supplemented Leibovitz’s L-15 Medium and detached cells were distributed into sterile flasks with fresh media. PAC2 cells were obtained from the Antonellis Lab at the University of Michigan (RRID: CVCL_5853).

### Electroporation and luciferase reporter assay

To assess the activity of the CMV enhancer in zebrafish cell culture, 4000ng of firefly luciferase expressing plasmids either with or without the CMV enhancer were transfected into 2×10^6^ PAC2 cells via electroporation (**Fig. 1A**). 100ng of the renilla luciferase expressing plasmid (pRL-SV40 Promega, E2231) was included as a transfection control. Firefly/renilla luciferase signal was calculated as the mean of ratios of three technical replicates per biological replicate. Fold change was then calculated relative to the signal of the SV40 promoter-only containing plasmid. The mean of fold-changes is reported and error bars represent standard deviation.

To test the functionality of our dual module *lac* repressor system in zebrafish cell culture, 2000ng repressible module and 2000ng of LacI-expressing plasmid were co-transfected into 2×10^6^ PAC2 cells by electroporation. 400ng of pRL-SV40 was included as a transfection control (**Fig. 2**).

All transfections were completed using 2mM cuvettes (Bulldog Bio, 12358-346) and electroporated using a NEPA21 Electroporator (Nepagene). Cells were harvested from culture and resuspended in 90μL of Opti-MEM Reduced Serum Medium (ThermoFisher, 31985062) per 1×10^6^ cells. Mastermixes of cells and DNA were prepared according to scale of conditions, and distributed into cuvettes (100μL/cuvette, 10μL of DNA and 90μL of cells). Poring pulse for PAC2 cells was set to the following: 200V, Length 5ms, Interval 50ms, Number of pulses 2, D rate% 10, Polarity +. Poring pulse for K562 was set to the following: 275V, Length 5ms, Interval 50 ms, Number of pulses 1, D rate 10%, Polarity +. For both cell types, the transfer pulse conditions were set to the following: 20V, Pulse length 50ms, Pulse interval 50ms, number of pulses 5, D rate 40%, Polarity +/-. Immediately following electroporation, each cuvette was recovered in 900μL of appropriate media and distributed into a well on a 24-well culture plate. For the transfection in Figure 2, PAC2 cells were recovered in 6-well plates. Each condition had a total of 3 biological replicates. For experiments including LacI, IPTG was treated as a separate condition and added to 1mM final concentration, unless otherwise specified, at 1 hour and 24 hours post-transfection (**Sup. Fig. 1**).

Luciferase results were collected 48 hours post-transfection on a GloMax-Multi+ Detection System (Promega, E7081) using the Promega Dual-Glo Luciferase Assay System (Promega, E2940).

### Zebrafish microinjections

Microinjections were carried out using the *Tol2* transposon system as previously described [24, 28]. Zebrafish embryos were co-injected within 30 minutes of fertilization in the single-cell stage with *Tol2* mRNA, the experimental plasmid, and phenol red for visualization. All embryos were maintained in 1X Holt buffer and fluorescent activity assessed at 24, 48, and 72 hours post-fertilization.

## Declarations

### Ethics Approval and consent to participate

This research was approved by the University of Michigan’s University Committee on Use and Care of Animals (protocol number PRO00008385).

### Consent for Publication

Not applicable

## Availability of Data and Materials

Data sharing is not applicable to this article as no datasets were generated or analysed during the current study.

## Author’s Contributions

SN performed and analyzed the CMV validation in cells and whole fish. TM performed and analyzed the *lac* operator-repressor system validations in cells. GF and MH assisted in the *Iac* operator-repressor system experiments. MD and JS generated key components necessary to the completion of these experiments and substantially contributed to the design of this work. AB made substantial contributions toward the conception of this work. SN, TM, GF, MD, JS, and AB wrote the manuscript. All authors read and approved the final manuscript.

## Acknowledgements

We thank Dr. Anthony Antonellis for the use of his zebrafish facilities, Dr. John Moran for the use of his plate reader, and Dr. Shigeki Iwase for the use of his Nepagene21 electroporator system.

## Supplemental Material

### Supplemental Methods

#### Cell Culture

The human myelogenous leukemia cell line K562 was cultured in RPMI 1640 + glutamine (ThermoFisher, 11875093) supplemented with 10% heat inactivated fetal bovine serum and 1% antibiotic-antimycotic until confluent. During growth K562 were incubated at 37°C in 5% CO2. Cells were split every two days into fresh media to avoid extended confluence.

#### Electroporation and Luciferase Reporter Assay

To determine the optimal amount of IPTG, we co-transfected 400ng of repressible reporter plasmid and 400ng of LacI-expressing plasmid into 1×10^6^ K562 cells by electroporation. We recovered cells in supplemented media containing 0, 1, 2.5, 5, 10, 20, or 50mM of IPTG. We included 75ng of Renilla luciferase expressing plasmid (pRL-SV40) as a transfection control (**Sup. Fig. 1**).

To compare the LacI dual module between different human and zebrafish cell types, we co-transfected 500ng of repressible module with 500ng of LacI-expressing plasmids into 1×10^6^ K562 cells or PAC2 cells by electroporation. 100ng of pRL-SV40 plasmid was included as a transfection control (**Sup. Fig. 2**).

## Supplemental Figures

**Supplemental Figure 1.**
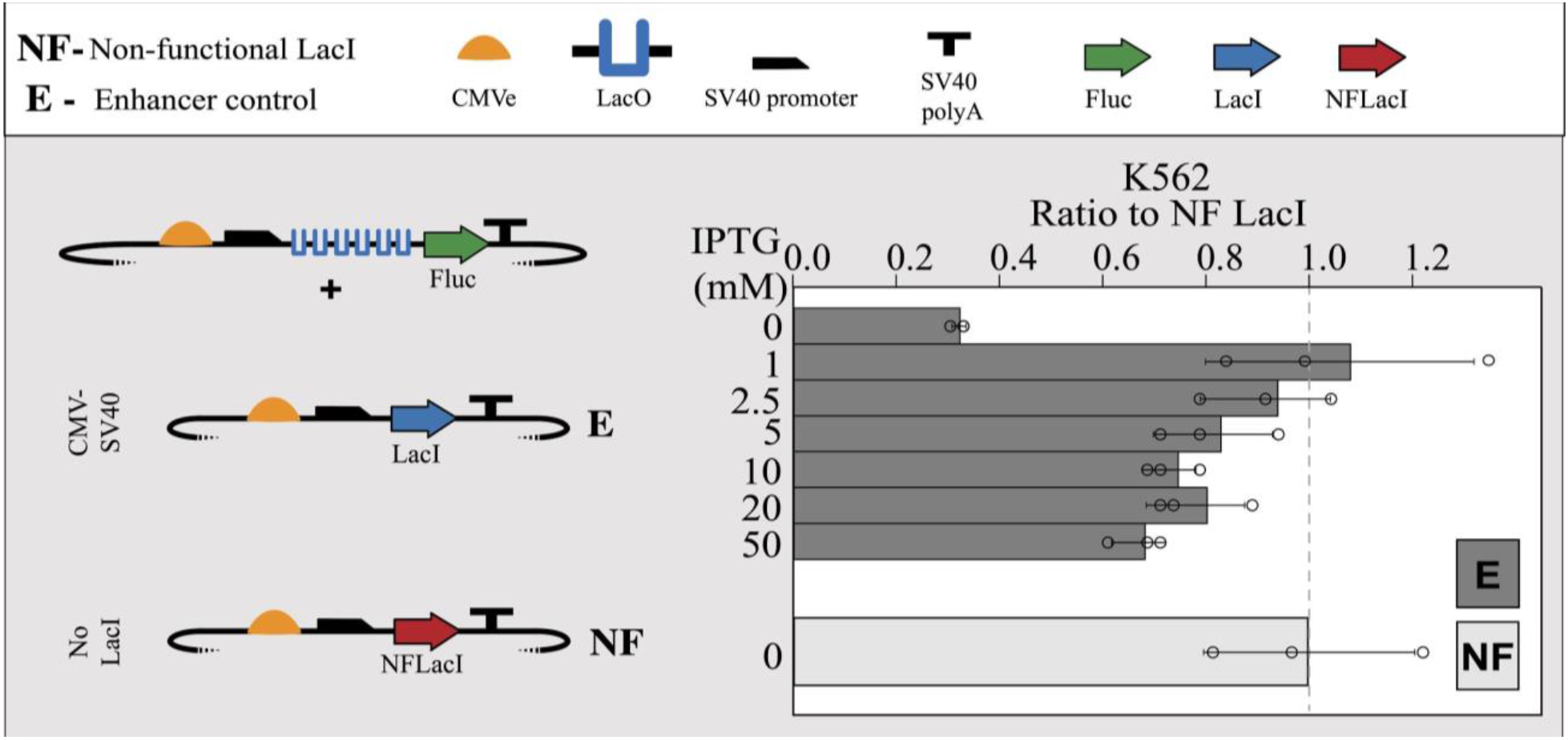
High levels of IPTG negatively impact the output of the *lac* operator-repressor system in K562 cells. Co-transfections of repressible reporter plasmids with equimolar amounts of CMV-SV40 enhancer-promoter enhancer driven LacI-expressing plasmids were exposed to increasing levels of IPTG. At 1mM IPTG, the output signal is restored to the level observed when NFLacI-expressing plasmids are co-transfected with repressible reporter plasmids, indicating that 1mM IPTG is sufficient to relieve LacI repression. At IPTG concentrations >1mM an increasingly lower output of signal relative to NFLacI levels was observed. The TSS begins where the SV40 promoter begins to slope downward. Error bars represent standard deviation of replicates (n=3). Points represent values for all 3 replicates in each condition. The dashed line shows the NF LacI IPTG-negative control.

**Supplemental Figure 2.**
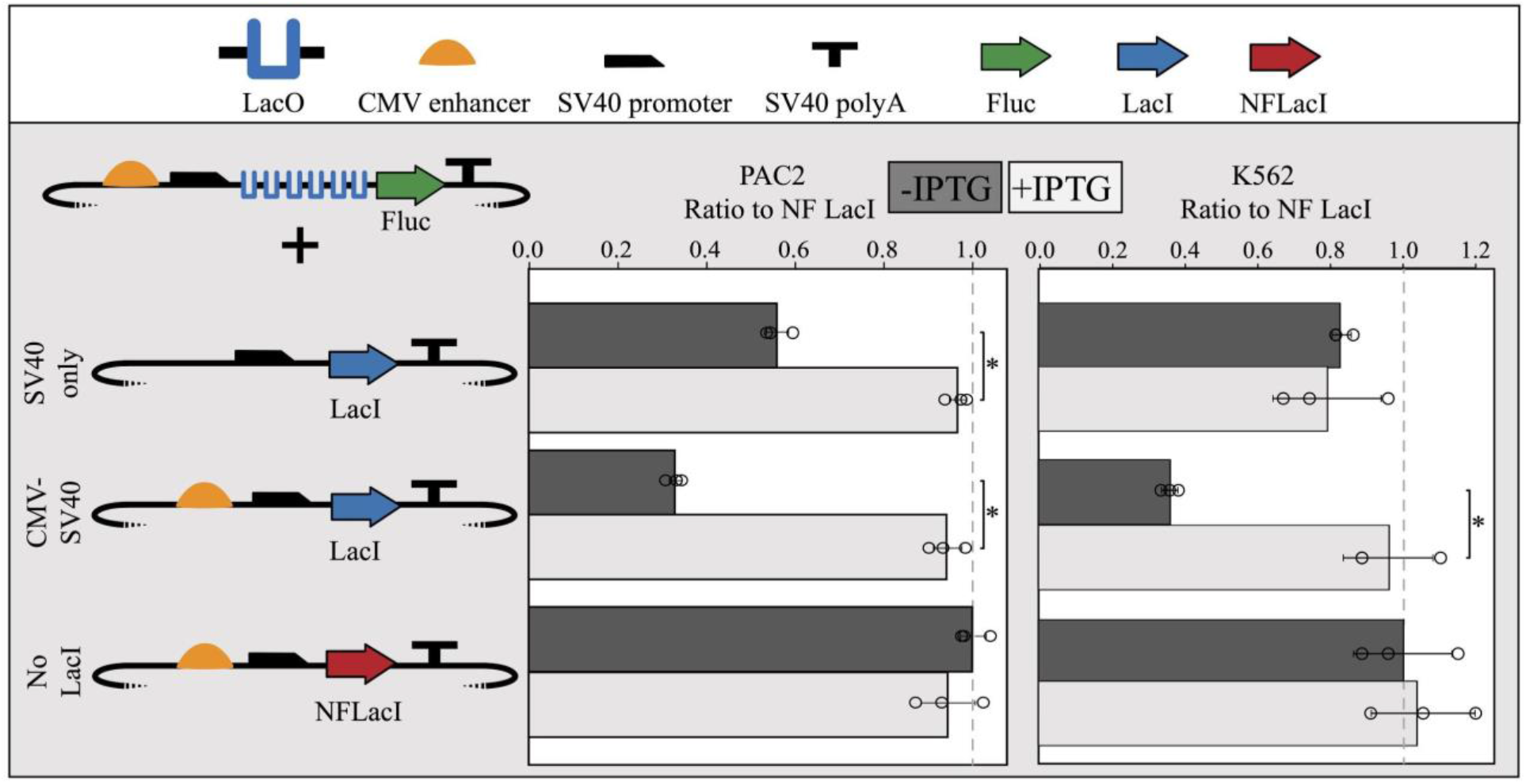
The *lac* operator-repressor system performs similarly in both human and zebrafish cell lines. K562 and PAC2 cells transfected with the same *lac* operator-repressor plasmid mixtures result in similar repression profiles. The promoter-only LacI-expressing plasmids resulted in roughly 10-20% repression in both cell types and the CMV-SV40 enhancer-promoter driven LacI-expressing plasmids resulted in ∼60% repression in both cell types. For each of the 3 plasmid combinations above, 100ng of pRL, 4:1 molar equivalents of repressible module plasmid:pRL, and 4:1 molar equivalents of LacI-expressing plasmid:pRL were co-transfected into 1 million K562 cells or PAC2 cells. 6 biological replicates were performed for each condition and 3 were exposed to 1mM IPTG and the remaining 3 were not exposed to IPTG. The TSS begins where the SV40 promoter begins to slope downward. Error bars represent standard deviation of replicates (n=3). Points represent values for all 3 replicates in each condition. The dashed line shows the NF LacI IPTG-negative control.

